# No One Is Leaving This Time: Appraisal of Awareness and Knowledge of Port Health Officers on International Health Regulation IHR (2005)

**DOI:** 10.1101/2022.01.12.476011

**Authors:** Yusuf Babatunde Adiama, Solomon Olayinka Adewoye, Opasola Afolabi Olaniyi, Habeeb Modupe Lateefat, Abdullahi Ahmed, Morufu Olalekan Raimi

**Affiliations:** Department of Environmental Health, Faculty of Pure and Applied Science, Kwara State University, Malete, Kwara State. Nigeria; Department of Epidemiology and Community Medicine, Federal Medical Centre Abeokuta, Ogun State, Nigeria; Department of Community Medicine, Environmental Health Unit, Faculty of Clinical Sciences, Niger Delta University, Wilberforce Island, Bayelsa State, Nigeria

**Keywords:** Seaport, International Health Regulation (IHR), Core Public Health Capacity, Awareness and Knowledge

## Abstract

**Background:** Historically, ships have played an important role in transmitting infectious diseases around the world. The spread of cholera pandemics in the 19^th^ century was thought to be linked to trade routes and facilitated by merchant shipping. The international maritime traffic of people and goods has often contributed to the spread of pathogens affecting public health.

**Objectives:** To assess level of awareness and knowledge of international Health regulation (IHR 2005) content among port health officer

**Methods:** The study design was descriptive cross-sectional evaluation, questionnaires were used to capture the respondents’ knowledge, awareness and sanitary condition of ship in accordance with (IHR 2005)

**Results:** On awareness and knowledge, Majority of the respondent (77.1 %) demonstrate good awareness of the IHR (2005), while 22.9% had not and some even testified of hearing the said document for the first time. Despite the fact that majority of respondent were aware but only 24.6% of them can actually demonstrate good knowledge of IHR (2005) and its intent to protect and prevent spread of disease along the international route.

**Conclusion:** There is need to improve the knowledge of port health officers by expand training and guidance on application of the IHR’s to frontline officer at point of entries. Also ensure more thorough inspection and avoid influence of ship agent during inspection of ship.

## 1. Introduction

Historically, ships have played an important role in transmitting infectious diseases around the world. The spread of cholera pandemics in the 19th century was thought to be linked to trade routes and facilitated by merchant shipping [1, 2]. According to Katz and Fischer [3] in the 1800s, the global community recognized the potential spread of diseases (particularly cholera, plague and yellow fever) across international borders [4, 5]. Quarantine was used to prevent the spread of these diseases across international borders and this brought about the implementation of the International Health Regulations (IHR). The International Sanitary Regulations were first adopted in 1951. In 1969, they were renamed as the International Health Regulations (IHR). The 1951 IHR were intended to monitor and control six serious infectious diseases: cholera, plague, yellow fever, smallpox, relapsing fever and typhus. In the intervening 50 years, many developments affected the international transmission of disease, including changes in international ship traffic. Therefore, on 23 May 2005, the World Health Assembly adopted a revised IHR by way of resolution WHA58.3, which entered into force on 15 June 2007. At the same time, the IHR clarifies a series of procedures that should be observed by the WHO to protect global public health safety [6]. The revised IHR focuses on public health crisis prevention, which has been expanded from certain “quarantine diseases” to any public health emergencies that may cause international repercussions. The implementation of the IHR shifts from the passive barrier of entry and exit points to the proactive risk management, aiming at early detection of any international threat before its formation and at stopping it from the very beginning [7].

## 2 Knowledge and Awareness of IHR 2005

Awareness of IHR [7] among port health staff was low. The staff has not been given any training on IHR related activities, and an assessment of training needs had not been carried out during the last five years. Only 1 port health officer (PHO) has undergone foreign training twice (on issuing ship sanitation certificates regarding SARS), but it was not related to IHR [7]. However, the training programs for the Ministry of Health (MOH) conducted omits the IHR [7].

Numerous researchers like Olalekan [8], Olalekan *et al*., [9], IMO [10]. Bakari and Frumence [11] reported low level knowledge of IHR (2005) among health workers in their studies; Some respondents had the correct understanding of IHR requirements. However, some had little information about the IHR [7]. It was noted that there was no training programme for Julius Nyere International Airport (JNIA) health workers for career development, knowledge and skills update. One respondent complained that she had not attended any training after her basic education. Lack of awareness, advocacy and adequate training on the importance of point of entry (POE) in implementing IHR [7] has left behind the efforts to spearhead the required strategies for disease surveillance and response systems [6]. Training and re-training of health workers at POE is part of preparedness and response, and therefore, it is crucial for JNIA health workers for career development and update of knowledge and skill, particularly on case detection. Opportunities for continuing education and upgrading skills should be initiated, particularly technical and managerial skills to prevent and control infectious diseases at JNI [12]. An assessment of human resource capacities and corresponding training needs in light of the IHR [7] multi-hazards approach was not undertaken. There were at least 17 health training institutions in the country for training public health specialists/epidemiologists, clinical medicine specialists, medical doctors, clinical officers, nurses and other paramedical professions [8, 9]. However, there were no training programs in epidemiology for diploma holders [13]. In a study done on training personnel on hygiene inspection on passenger ships, 17 authorities (39.5%) have received training on hygiene inspections, while 11.1% have received training specifically for ship inspections. The majority of the responding authorities (73.2%) believe that specific training for passenger ship inspections is needed [14].

Analysis of results on funding sources for training workforce shows that even when such funding was provided, it was usually grossly inadequate to serve any meaningful purpose as it concerns implementing the IHR requirements. For example, respondents reported that the usual practice was for the various POE port health services to send work plans containing funding requirements to the government through the ministry of health. As a result, human resources training were inadequate across the POEs. On the frequency of the international training, 82.9% of the respondent at the seaport session reported having not attended any training. However, appreciable levels of well-trained personnel were documented at the sea and airport, respectively. About 70% of the respondents across all categories in this study said they had not attended any training on the content of IHR requirements [15]. This confirmed Uganda study on training needs considering the IHR [7] multi-hazards approach had not been undertaken. Several studies have reported the need for countries to identify and mobilize the required technical, financial and human resources from all possible available sources to focus on the implementation of IHR [7] to meet core capacity requirements [16]. Muhammed *et al*. [15] reported that knowledge and training need shows 26.8% of the respondents at the Airport are trained on the content of IHR, while 73.2% of the respondents are not trained on the content of IHR. Capacity developments in terms of training of core personnel on the content of IHR 2005 across the three. A total of 37 articles from 49 countries reported experience implementing the human resources core capacity from Cambodia, India, Uganda, and the United Kingdom indicated that traditional curricula, competencies and training did not prepare the workforce to implement IHR [7] and that additional knowledge transfer and skill-building is needed to ensure reporting and data use at subnational levels [17]. They also reviewed that sociocultural context influenced learning preferences varies from country to country and reported that face-to-face learning was preferred in Morocco, while electronic learning was preferred in India and the United Kingdom. In China, Morocco, Africa and the United Kingdom, interactive and skill-building sessions were preferred over static knowledge transfer. Settings that had high staff turnover (e.g. rural areas, those with armed conflicts) faced staff shortages and required unique mechanisms for continual retraining.

## 3 Methodology

The study was conducted in some selected seaports in Nigeria. The Federal Republic of Nigeria is in West Africa on the Gulf of Guinea coast. The study design was descriptive cross-sectional evaluation. Questionnaires were used to capture the respondents’ knowledge, awareness, experiences, and their understanding of IHR [7] and its content, in line with Core capacity requirements raised in the IHR [7] for human resource for health. The study covered Port health officers (PHOs) in Apapa Sea port, Warri and Port Harcourt seaports. Port health officer working in selected Seaports with sufficient cognitive ability to answer the study questions that were included.

### Sample Size Determination

- The minimum sample size was determined using the Fischer’s formula for obtaining sample size
- size when the population is less than 10,000 for descriptive studies

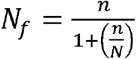

Value of n was calculated using the formula n = <u>Z^2^ pq</u>

d^2^Araoye, [18].

q = 1.0 – p

d = degree of accuracy, set at 0.05 for this study.

- Adjustment for Non-Response
- The minimum sample size; N = n/(100-r %)

Where r% was the anticipated non response rate, which is 10%

- A total of 179 respondents (each for PHO and ships for inspection) were used for this study.

A systematic sampling technique was used for the selection of respondents.

All the selected site were used for the study. The numbers of PHOs interviewed were proportionately allocated based on the number of PHO in each selected sea port and equal allocation distribution were done for the ships.

- Allocation at each port level was based on the sampling-by-size approach, using the formula below:

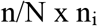

### Ethical statements

Verbal and written consent was obtained from every respondent after explaining the study’s purpose and assured the confidentiality of information obtained from them. Ethical clearance was obtained from KWASU Institutional Research Board and Publication Committee. Thus, ethical clearance for the study was obtained from the Research Ethics Committee of Kwara state University, Malete-Nigeria (Reference number: KWASU/FPAS/EHS/023). Verbal and written consent were sought from the respondents after explaining the study details, and importance of the research work. The respondents were assured the personal information will be treated with utmost confidentiality.

### Statistical analysis

The administered questionnaires were analyzed using appropriate statistical analyses such as frequencies, Chi-square test, and Ordinal logistics regression.

## 4. Results

Table 1 shows the modal age group is 44 – 47yrs (34.9%). Majority of the respondents were male, constituting 88.9%, 61.9% and 84.1 across the selected seaport. 158 (88.3%) were married while 6 (3.4%).5% were singled. The majority 151 (84.4%%) of the respondents, had Bsc/Bed/HND as their highest level of education but none with Ph.D. Islam is the religion practiced by 54.7% of the respondents.

**Table 1:**
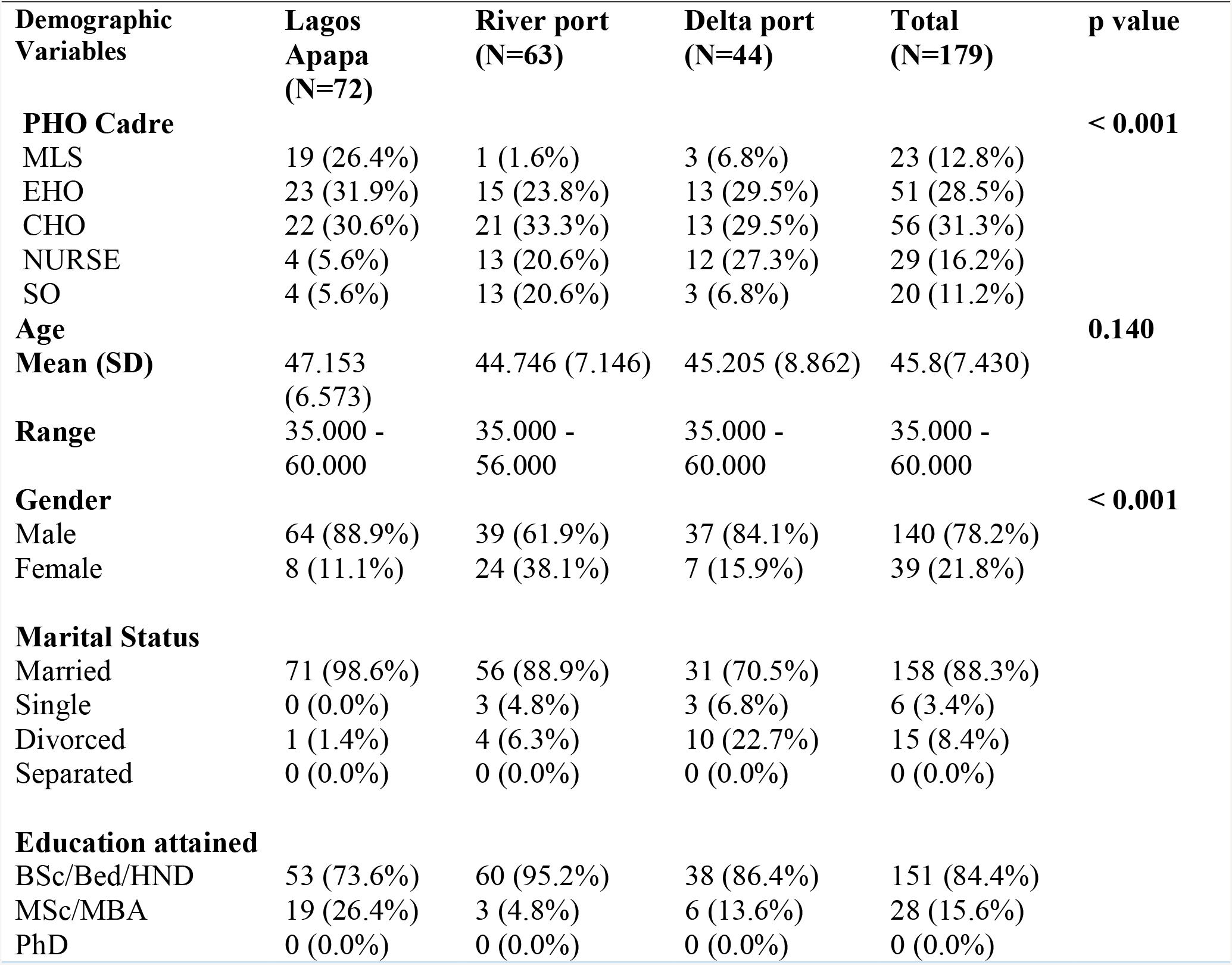
Socio-Demographic Characteristics of Respondents.

| **Demographic Variables** | **Lagos Apapa (N=72)** | **River port (N=63)** | **Delta port (N=44)** | **Total (N=179)** | **p value** |
| --- | --- | --- | --- | --- | --- |
| **PHO Cadre** |  |  |  |  | **< 0.001** |
| MLS | 19 (26.4%) | 1 (1.6%) | 3 (6.8%) | 23 (12.8%) |  |
| EHO | 23 (31.9%) | 15 (23.8%) | 13 (29.5%) | 51 (28.5%) |  |
| CHO | 22 (30.6%) | 21 (33.3%) | 13 (29.5%) | 56 (31.3%) |  |
| NURSE | 4 (5.6%) | 13 (20.6%) | 12 (27.3%) | 29 (16.2%) |  |
| SO | 4 (5.6%) | 13 (20.6%) | 3 (6.8%) | 20 (11.2%) |  |
| **Age** |  |  |  |  | **0.140** |
| **Mean (SD)** | 47.153 (6.573) | 44.746 (7.146) | 45.205 (8.862) | 45.8(7.430) |  |
| **Range** | 35.000 - 60.000 | 35.000 - 56.000 | 35.000 - 60.000 | 35.000 - 60.000 |  |
| **Gender** |  |  |  |  | **< 0.001** |
| Male | 64 (88.9%) | 39 (61.9%) | 37 (84.1%) | 140 (78.2%) |  |
| Female | 8 (11.1%) | 24 (38.1%) | 7 (15.9%) | 39 (21.8%) |  |
| **Marital Status** |  |  |  |  |  |
| Married | 71 (98.6%) | 56 (88.9%) | 31 (70.5%) | 158 (88.3%) |  |
| Single | 0 (0.0%) | 3 (4.8%) | 3 (6.8%) | 6 (3.4%) |  |
| Divorced | 1 (1.4%) | 4 (6.3%) | 10 (22.7%) | 15 (8.4%) |  |
| Separated | 0 (0.0%) | 0 (0.0%) | 0 (0.0%) | 0 (0.0%) |  |
| **Education attained** |  |  |  |  |  |
| BSc/Bed/HND | 53 (73.6%) | 60 (95.2%) | 38 (86.4%) | 151 (84.4%) |  |
| MSc/MBA | 19 (26.4%) | 3 (4.8%) | 6 (13.6%) | 28 (15.6%) |  |
| PhD | 0 (0.0%) | 0 (0.0%) | 0 (0.0%) | 0 (0.0%) |  |

**Table 2:** Awareness and Knowledge of the Respondents on IHR (2005)

|  |  | **Lagos Apapa** | **River port** | **Delta port** |
| --- | --- | --- | --- | --- |
|  |  | **(n=72)** | **(n=63)** | **(n=44)** |
| Are you aware of the existence of a document called IHR | Yes | 67(93.1%) | 35(55.6%) | 36(81.8%) |
|  | No | 5(6.9%) | 28(44.4%) | 8(18.2%) |
| Do you know its objectives | Yes | 22(30.6%) | 16(25.4%) | 24(54.5%) |
|  | No | 50(69.4%) | 47(74.6%) | 20(45.5%) |
| Knowledge on core capacity Requirement | Yes | 31(43.1%) | 7(11.1%) | 26(59.1%) |
|  | No | 41(56.9%) | 56(88.9%) | 18(40.9%) |
| Do you have a copy of IHR | Yes | 21(29.2%) | 3(4.8%) | - |
|  | No | 51(70.8%) | 60(95.2%) |  |
| Training on IHR | Yes | 22(30.6%) | 10(15.9%) | 4(9.1%) |
|  | No | 50(69.4%) | 58(92.1%) | 40(90.9%) |

Majority of respondents, 67(93.1%), 36(81.8%), 35(55.5%) at Apapa, River and Delta seaport, are aware of IHR [7]. While 24(54.5%), 22(30.6%) of Lagos and Delta port know its objectives and intent to prevent diseases at the point of entry. 22(30.6%) of respondents from Lagos port had attended training on IHR and 4(9.1%) from Delta.

Figure 1 shows that the majority of 89(49.0%) of the respondents had on-the-job training, while 60 (33.5%) of respondents had face-face training courses.

**Figure 1:**
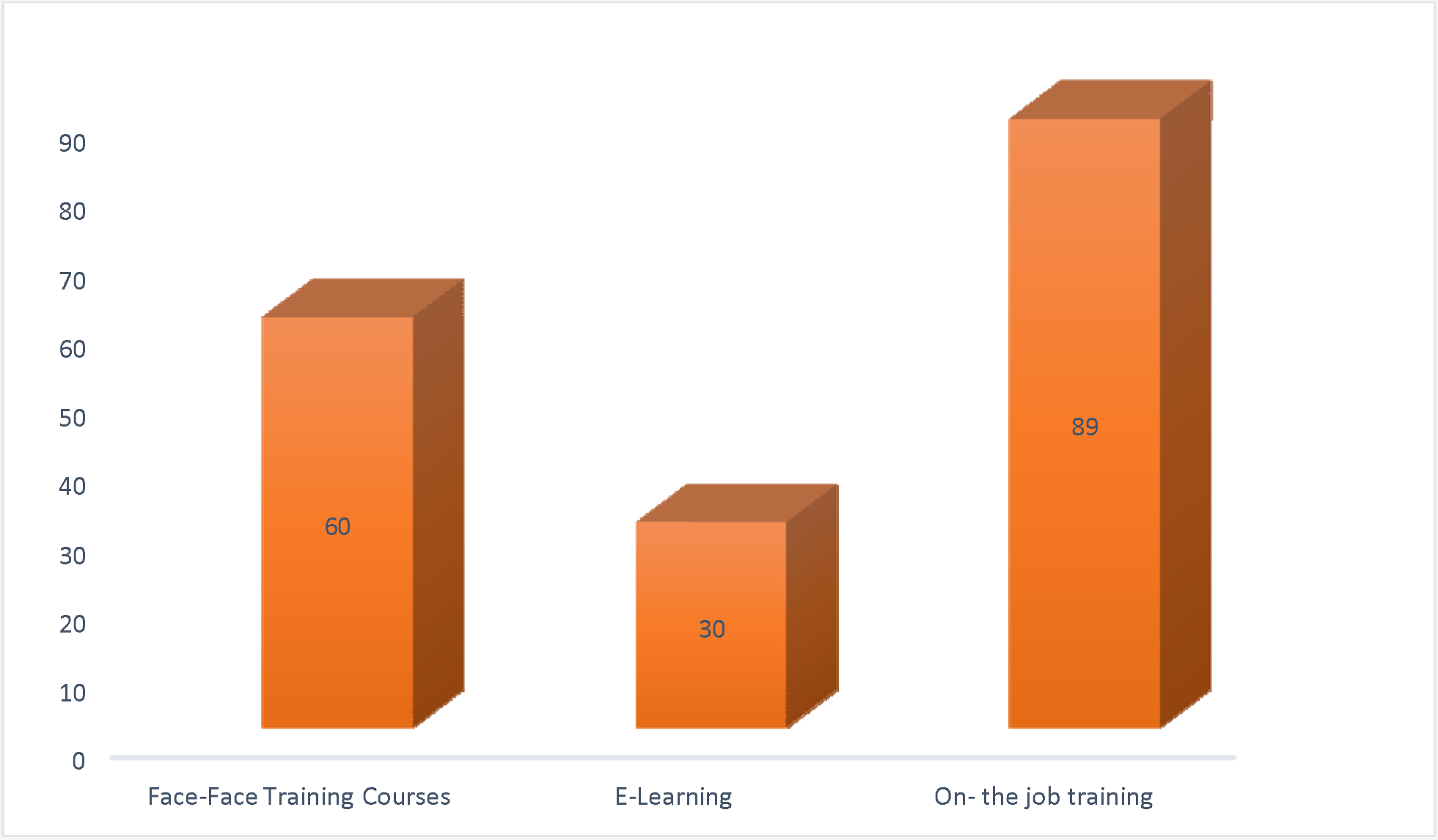
Mode of training by the respondents

Figure 2 shows that on the training frequency, only 15 (8.3%) of the respondents said the training takes place on an annual basis while the majority of respondents, 139 (77.7%), responded as the case may be and that no definite time and period for training.

**Figure 2:**
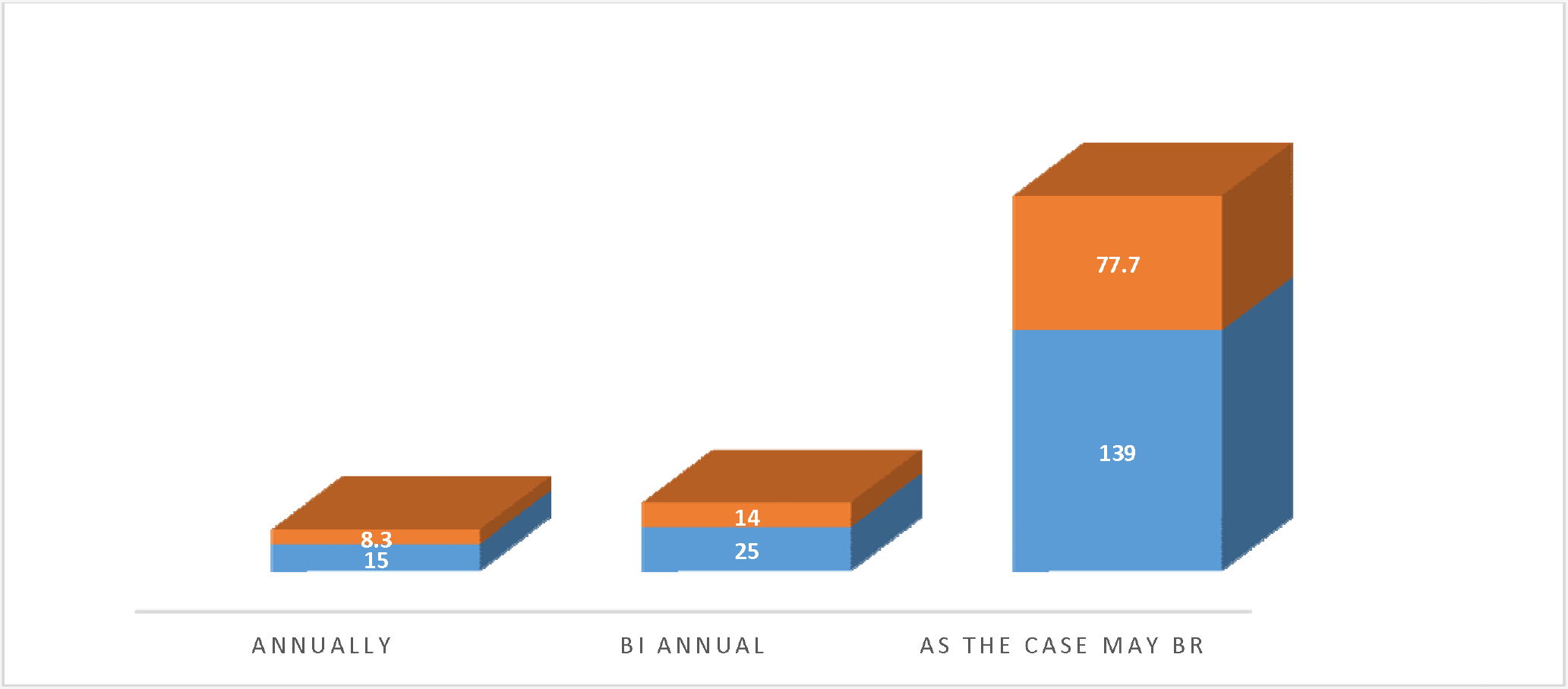
Frequency of training by the respondents

From Table 3 and figure 3 shows the p-values (0.02351) is less than the level of significance(α = 0.05), this indicates a strong association between the awareness and knowledge of international health regulations [7] with various departments/units. The plots below also buttress the awareness and knowledge of international health regulations [7] with various departments/units.

**Table 3:** Relationship of Awareness and Knowledge with Various Department/Cadre

| Department/Cadre | Awareness | | $\chi^2$ Statistics | Degree of freedom (df) | P-value |
| --- | --- | --- | --- | --- | --- |
|  | Yes | No |  |  |  |
| CHO | 30 | 8 | 11.288 | 4 | 0.02351 |
| EHO | 72 | 10 |  |  |  |
| Doctor | 7 | 3 |  |  |  |
| Nurse | 4 | 6 |  |  |  |
| Sc. Office | 9 | 4 |  |  |  |

**Figure 3:**
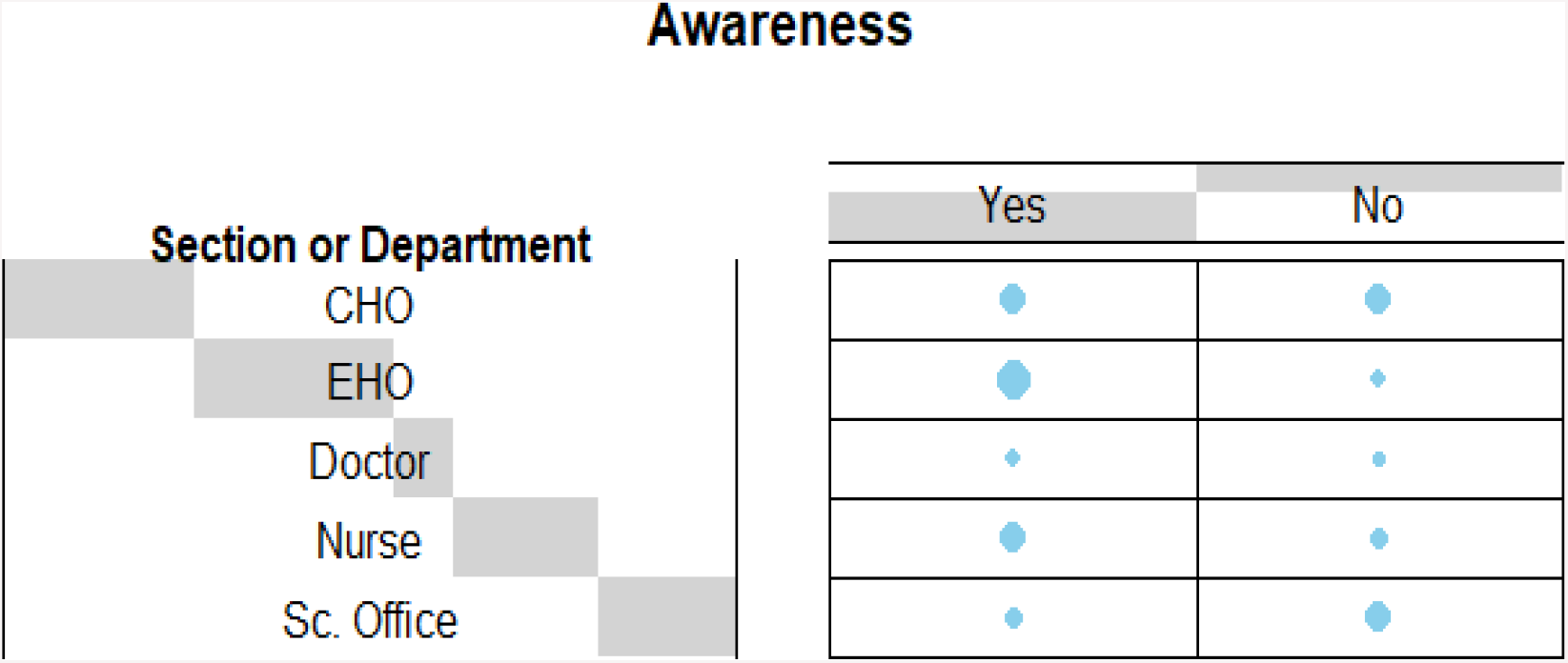
Awareness with Various Department/Cadre

Figure 4 shows the rate of awareness and knowledge across the seaport by ordinal logistic regression with the area under curve 0.848, 0.843and 0.755 shows Lagos, River and Delta port, respectively. Moreover, this indicated Lagos seaport is more aware and have good knowledge when compared with other seaports in the study sites.

**Figure 4:**
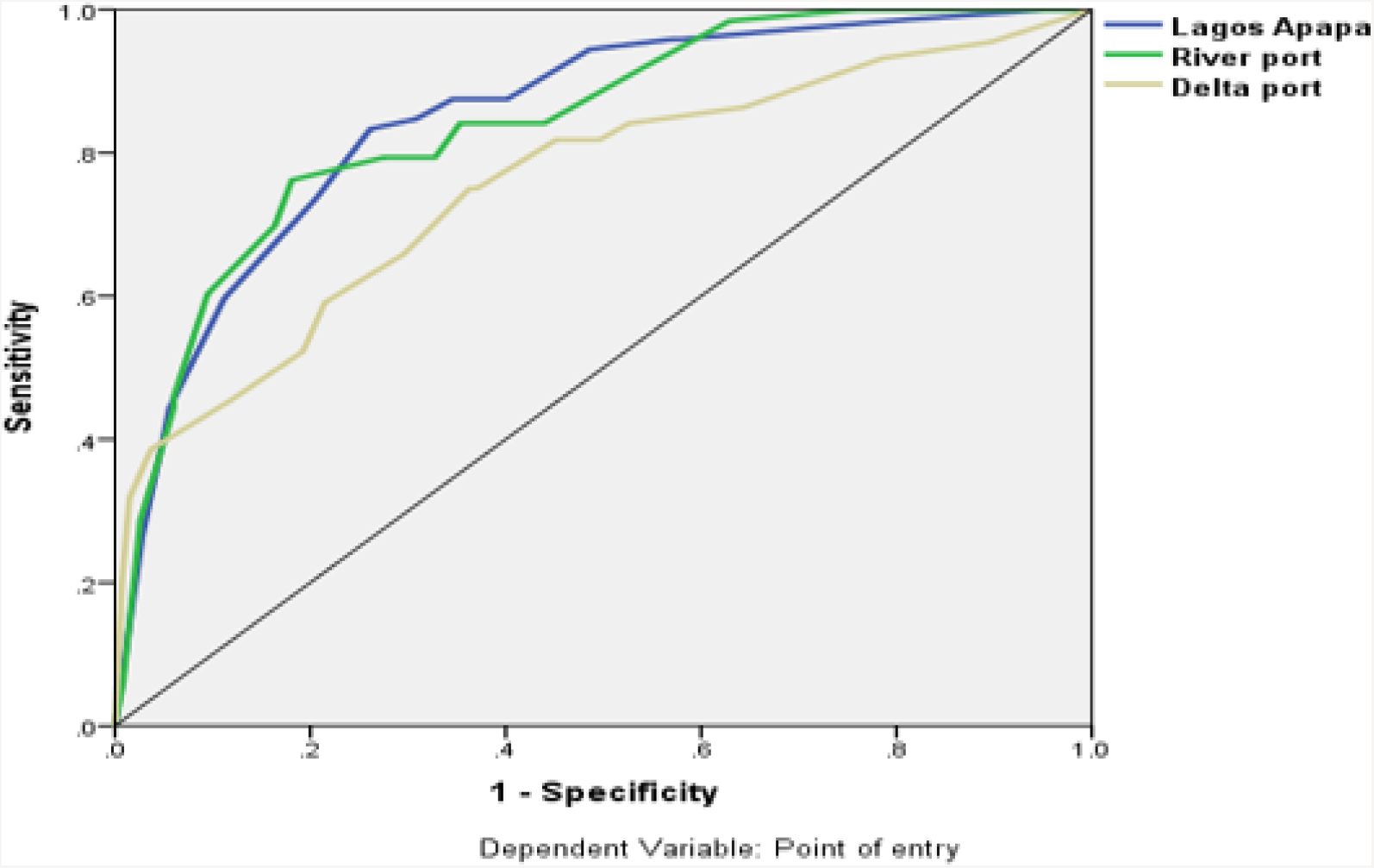
Rate of awareness and knowledge of IHR [7]

Table 4 shows that the majority, 138 (77.1%) of respondents across the selected seaports, had good awareness, but only 44(24.6) had good knowledge about IHR (2005)

**Table 4:** Awareness and Knowledge Score of the Respondents on IHR (2005) N= 179.

| Variable | Score | Frequency | Percent |
| --- | --- | --- | --- |
| Awareness | Good | 138 | 77.1 |
|  | Poor | 41 | 22.9 |
| Knowledge | Good | 44 | 24.6 |
|  | Poor | 135 | 75.4 |

## 5 Discussion of the Findings

### Awareness and Knowledge of Respondents

On awareness and knowledge, majority of the respondents (77.1%) demonstrate a good awareness of the IHR [7], while 22.9% had not and some even testified of hearing the said document for the first time. Even though most respondents are aware, only 24.6% of them can actually demonstrate good knowledge of IHR [7] and its intent to protect and prevent the spread of disease along the international route. Since the P-values (0.02351) is less than the level of significance(α = 0.05), this indicates a strong association between the awareness and knowledge of international health regulations [7] with various departments/units. On the rate of awareness and knowledge of IHR [7] across the selected seaport ordinal logistic regression was used for precision accuracy and revealed that Apapa Seaport (0.848) followed River seaport (0.0845) and Warri seaport (0.755) in Delta State. This finding was dissimilar to the study conducted by IMO (2013), where awareness of IHR [7] among port health staff was low. Also, the study confirmed Bakari and Frumence [11] assertion on low level knowledge of IHR [7] among health workers in his study. Some respondents had the correct understanding of IHR requirements. This finding was dissimilar to the study conducted by IMO [10], where awareness of IHR [7] among port health staff was low. Also, the study confirmed Bakari and Frumence [11] assertion on low level knowledge of IHR (2005) among health workers in his study. Anema *et*.*al* [19] reported in his study conducted among National focal person on IHR revealed, 88% reported having excellent/good knowledge of IHR [7] Annex 2; and this was contrary to this study 75.4% reported poor knowledge of it assessing potential public health emergency of international concerns (PHEICs). However, some of them had little information about the IHR (2005). Some respondents had the correct understanding of IHR requirements. However, some had little information about the IHR [7]. On the issue of capacity development, especially on training to enhance knowledge, all respondents across the selected ports (30.6%, 15.9%, and 9.1%) revealed not to have had adequate training on IHR [7]. Muhammed *et al*. [15] affirmed this study, reported that knowledge and training need shows, 26.8% of the respondents at the Airport are trained on the content of IHR, while 73.2% of the respondents are not trained on the content of IHR [7] across the three points of entries. Amitabh *et al*. [17] showed experience from Cambodia, India, Uganda, and the United Kingdom indicated that traditional curricula, competencies and training did not prepare the workforce to implement IHR [7] and that additional knowledge transfer and skill-building are needed to ensure reporting and data use at the subnational level.

## 6 Conclusion

The need for knowledge and skills of related staff is important in border health especially port health officers who serve as gate way health officer in the process of developing, strengthening, and maintaining the core public health capacity both routine and emergency in order to prevent spread of disease of public emergency of international concern. Hence under scoring the need that Training efforts should be increased to build the legal and scientific sense to deal with health emergencies at the PoEs understudy. Frontline Health works at seaports are expected to be aware and know about basic tool i.e., IHR [7] of principle embedded. All theories, methods, and skills for processing and should apply them in practice in order for them to be able to implement the provisions of IHR 2005. Government on its part should continue its commendable efforts of health workforce development as noted in the report of the Joint External Evaluation, June 2017. This needs to be address in keeping with the global One Health approach mantra of not leaving no one behind this time.

## References

1. Cheesbrough J. S., Green J., Gallimore C. I., Wright P. A., and Brown D. W (2000). Widespread environmental contamination with Norwalk-like viruses (NLV) detected in a prolonged hotel outbreak of gastroenteritis. Epidemiology and Infection. 125(1):93–98. [PMC free article] [PubMed.

2. Raimi, M. O, Pigha Tarilayun K and Ochayi, E. O (2017) Water-Related Problems and Health Conditions in the Oil Producing Communities in Central Senatorial District of Bayelsa State. Imperial Journal of Interdisciplinary Research (IJIR) Vol-3, Issue-6, ISSN: 2454-1362.

3. Katz, R. & Fischer, J. (2011). The Revised International Health Regulations: A Framework for Global Pandemic Response. Global Health Governance, Spring 2010.3(2) http://www.ghgj.org (Retrieved on Dec 2nd 2011).

4. Morufu Olalekan Raimi, Tonye Vivien Odubo & Adedoyin Oluwatoyin Omidiji (2021) Creating the Healthiest Nation: Climate Change and Environmental Health Impacts in Nigeria: A Narrative Review. Scholink Sustainability in Environment. ISSN 2470-637X (Print) ISSN 2470-6388 (Online) Vol. 6, No. 1, 2021 www.scholink.org/ojs/index.php/se. URL: http://dx.doi.org/10.22158/se.v6n1p61. http://www.scholink.org/ojs/index.php/se/article/view/3684.

5. Raimi Morufu Olalekan, Tonye V. Odubo, Omidiji Adedoyin O, Oluwaseun E. Odipe (2018) Environmental Health and Climate Change in Nigeria. World Congress on Global Warming. Valencia, Spain. December 06-07, 2018

6. Chretien JP, Yingst SL, Thompson D (2010). Building public health capacity in Afghanistan to implement the International HealthRegulations: a role for security forces. Biosecurity Bioterror; 8: 277–85.

7. International Health Regulations (2005) International Health Regulations.Second edition. Available from: http://whqlibdoc.who.int/publications/2008/9789241580410_eng.pdf?ua=1

8. Olalekan RM (2020). “What we learn today is how we behave tomorrow”: a study on satisfaction level and implementation of environmental health ethics in Nigeria institutions. Open Access Journal of Science; 4(3):82lJ92. DOI: 10.15406/oajs.2020.04.00156.

9. Olalekan RM, Olawale SH, Christian A, Simeon AO (2020). Practitioners Perspective of Ethical Cases and Policy Responses by Professional Regulator: The Case of Environmental Health Officers Registration Council of Nigeria (EHORECON). American Journal of Epidemiology & Public Health. 2020;4(1): 016–023. https://www.scireslit.com/PublicHealth/AJEPH-ID23.pdf https://www.scireslit.com/PublicHealth/articles.php?volume=4&issue=1

10. Resolution A.1087(28) Adopted on 4 December 2013 (Agenda item 12) 2013 Guidelines for the Designation of Special Areas Under Marpol.

11. Bakari, E., & Frumence, G. (2013) Challenges to the implementation of International Health Regulations (2005) on preventing infectious diseases: experience from Julius Nyerere International Airport, Tanzania. Glob Health Action. 2013;6(1):20942. doi: http://dx.doi.org/10.3402/gha.v6i0.20942.

12. World Health Organization (2011). Handbook for Inspection of Ships and Issuance of Ship Sanitation Certificates; WHO: Geneva, Switzerland, 2011.

13. Wamala, J.F., Okot, C., Makumbi, I., Natseri, N., Kisakya, A., & Nanyunja, M., et al. (2010). Assessment of Core Capacities for the International Health Regulations (IHR[2005])- Uganda, 2009. BioMed Central Public Health, 2010. 10(1).

14. Mouchtouri, V., Van Reusel, D., Bitsolas, N., Katsioulis, A., Van den Bogaert, R., Helewaut, B. & Hadjichristodoulou, C. (2018). European web-based platform for recording international health regulations ship sanitation certificates: Results and perspectives. International journal of environmental research and public health, 15(9), 1833.

15. Muhammad S.U., Oluwasogo A.O., Henry O. S (2018): Assessment of Human Resources Core Capacity under International Health Regulations 2005 (Ihr 2005) At Ports Of Entry (Poe) In Lagos. OSR Journal of Environmental Science, Toxicology and Food Technology (IOSR-JESTFT) e-ISSN: 2319-2402, p- ISSN: 2319-2399. Volume 12, Issue 9 Ver. II (September. 2018), PP 01-08 www.iosrjournals.org

16. Peralta SOP (2008). Challenges for IHR Implementation in Spain. Madrid: Ministry of Health and Consumers’ Affairs.

17. Amitabh, B., Suthar, L.G., Allen, S.C., Christopher, D. & Jason, M.N. (2018). Lessons learnt from implementation of the international health regulations: a systematic review Article in Bulletin of the World Health Organization · February 2018.

18. Araoye, MO (2004). Sample size determination. Research Methodology with statistics for health and social sciences. Ilorin: Nathadex; p. 118–20.

19. Anema, A., Druyts, E., Hollmeyer, H.G., Hardiman, M.C., & Wilson, K. (2012). Descriptive review and evaluation of the functioning of the International Health Regulations (IHR) Annex 2. Vol. 8, Globalization and Health; 2012.

